# Fractions of traditionally brewed rice beverage ameliorate anxiety and improve spatial memory in mice

**DOI:** 10.1101/2021.05.01.442231

**Authors:** Bhuwan Bhaskar, Atanu Adak, Mojibur R. Khan

**Author notes:** Full list of author information is available at the end of the article.

## Abstract

Rice beverages are traditionally prepared and consumed popularly by the different ethnic groups of North East India. To investigate its effects on behavior, mice were treated with different fractions of rice beverage that included the beverage as a whole, insoluble and soluble fractions. Intragastric treatments of these fractions were given to the mice (n=6 per group) for 30 days and behavioral studies were performed on elevated plus and Y maze to evaluate anxiety and spatial memory, respectively. Next generation sequencing of metagenomic DNA of the beverage indicated presence of 157 OTUs and 26 bacterial genera were dominant with an abundance of 0.1%. The insoluble fraction treated animals showed lowest anxiety like symptoms. Spatial memory improved in all the treatments compared to the control, of which the rice beverage treatment showed the highest levels (*p*<0.05). Gas chromatography and mass spectroscopy-based metabolite profiling of the beverage revealed 10 alcohols, 29 sachharides, 43 acids and 13 amino acids. Findings of this study suggest a positive effect of rice beverage components on anxiety and spatial memory of mice.

## Introduction

North East India consists of eight states namely Arunachal Pradesh, Assam, Manipur, Meghalaya, Mizoram, Nagaland, Sikkim and Tripura **(Figure 1)** and phyysiographically categorized into the Eastern Himalayas, Northeast hills (Patkai-Naga Hills and Lushai Hills) and the Brahmaputra and Barak Valleys. The region is home to around 225 tribes which makes it rich in diverse ethnicity culture and food practices [1]. The foods prepared and consumed by these communities have a strong connection with their sociocultural, spiritual life and health. These foods include different types of fermented foods and beverages. The rich ethnic diversity and surplus availablity of bioresources contribute to a huge reserve of artisanal foods and beverages in this region which are often commercially available in small regional markets. Most of the ethnic communities of this zone prepare and consume fermented foods and beverages, however, the name, recipe and preparation method often varies tribe to tribe. Since rice is the major staple food of the region, a wide variety of rice based fermented alcoholic beverages are popularly consumed. Each tribe has a recipe for the preparation of the starter, which serves as the source of microbial inoculam and also the beverage has a different name based on the tribe which prepares it. Though the overall process remains same for each tribe, the differences lie in the compostion of starter, variety of rice to be used for fermentation and some additional steps during filtereation of the final product. The starter serves as inoculum consisting of starch degrading and alcohol producing microbes along with lactic acid bacteria. It has been reported to contain the bacteria *Lactobacillus plantarum, Lactobacillus brevis, Leuconostoc lactis, Weissella cibaria, Lactococcus lactis, Weissella para mesenteroides, Leuconostoc pseudomesenteroideetc* [2]. The starter is mixed with cooked rice under hygienic conditions for fermentation and the liquid is separated from the mass by decantation and filtration [3]. Consumption of rice beverage is traditionally believed to exert health benefits pertaining to improved digestive, excretory and endocrine functions and relieves stress [3]. Some of these benefits could be attributed to the presence of antioxidants, prebiotic components like raffinose, trehalose and probiotic bacteria along with alcohol, which can be beneficial in moderate amounts. The beverage has been reported to be rich in essential nutrients such as carbohydrates, amino acids, organic acids along with other aromatic compounds [4]. Sugars *viz*. mannose, trehalose and allose present in this beverage have prebiotic effect [5].The prebiotic and probiotic components confer health benefits like prevention of heart disease, diabetes, obesity and certain gastrointestinal diseases by stimulating the gut bacteria [6]. We hypothsize that the overall nutrients and microbes in the beverage may improve gut and mental function. This study aimed to explore the effects of rice beverage and its fractions on anxiety and spatial memory, using mouse model in an attempt to validate the claims by ethnic communities **(Figure 2)**.

**Figure 1.**
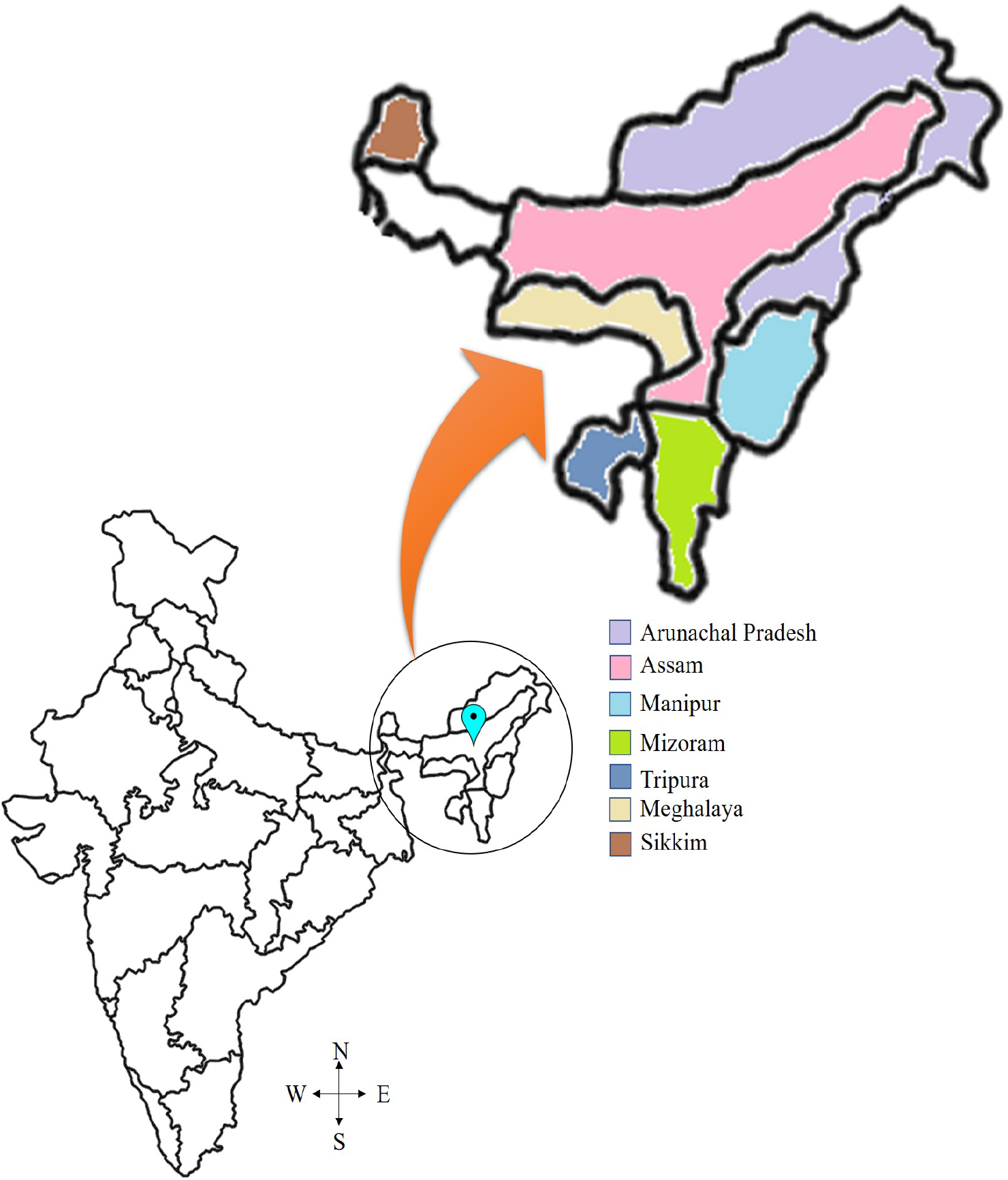
The seven states of North East India are inhabited by many indigenous communities having diverse food habits. Majority of these communities consume fermented foods and beverages. Assam is a state in this region with a rich diversity of rice based fermented beverages. The beverage holds an intrinsic value in the sociocultural life of these communities. (Map adapted from https://www.pikpng.com)

**Figure 2.**
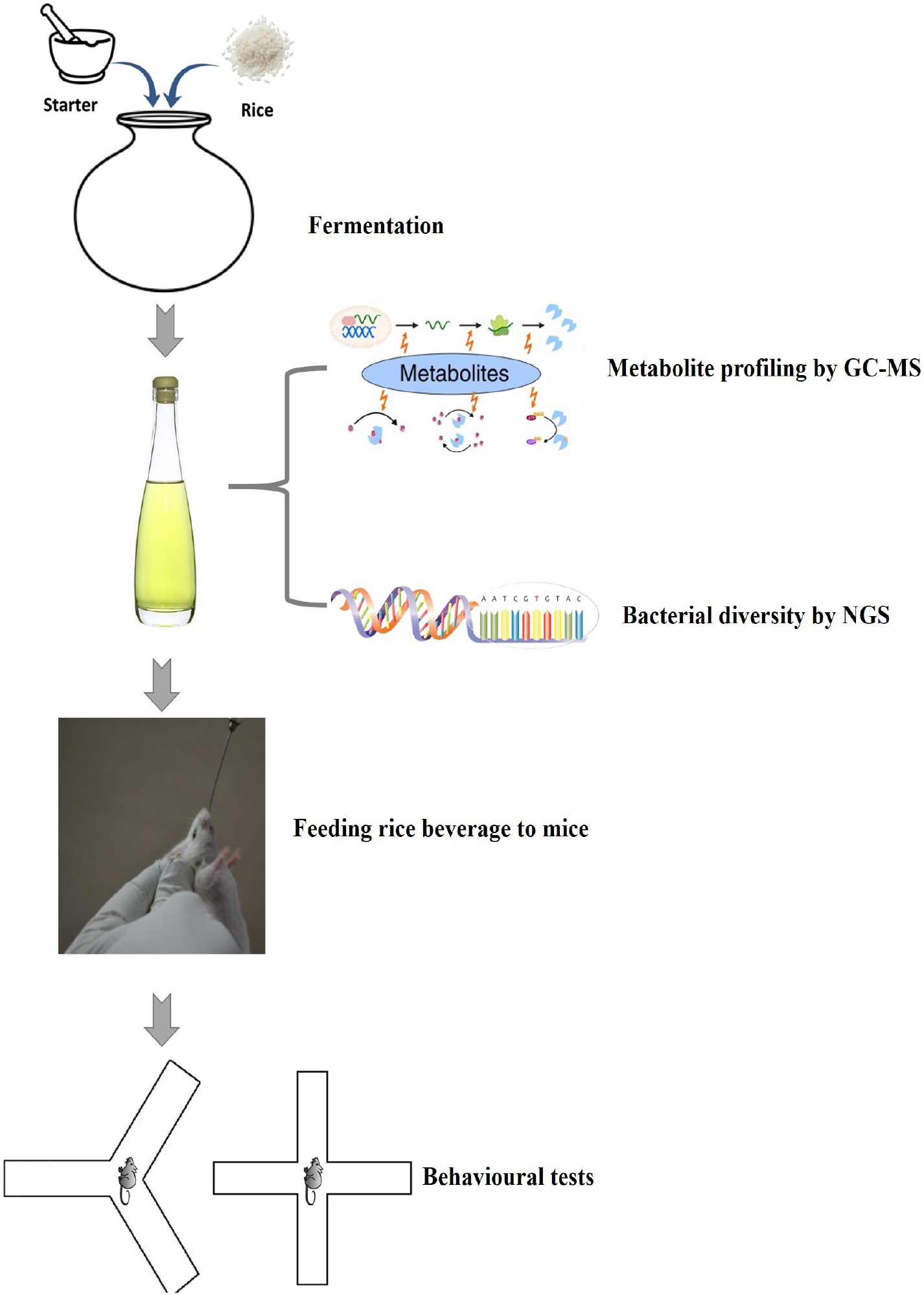
Summery of the experimental process performed in the study

## 1 Methods

### 1.1 Rice fermentation

Rice beverage was prepared based on the traditional knowledge of *Tiwa* community of Assam. The starter cake was prepared using different plants and rice as described in our earlier patent [7]. Further, 1 kg of rice *(Oryza sativa)* was boiled and allowed to cool down to room temperature. A 30 gm of starter cake, crushed into powder was mixed with the cooled rice and kept for solid state fermentation for 8 days at room temperature. After fermentation,the mass was sieved with a muslin fabric to obtain rice beverage (RB) which was further centrifuged at 5000 rpm for 5 min to separate the supernatant containing the soluble fraction (SF) and the pellet was resuspended in sterile water upto the initial volume, denoted as insoluble fraction (IF).

### 1.2 Biochemical analyses and non targeted profiling of rice beverage metabolites by Gas chromatography–mass spectrometry (GC-MS)

The pH was determined in a digital pH meter (pH510, Orion Star A111). The alcoholic fraction was evaporated and collected in a rotary evaporator to estimate the alcohol by volume using potassium dichromate reagent[8]. Reducing sugars, antioxidant activity and total phenolic content was determined as described elsewhere [9, 10, 11]. GC-MS was carried out as per protocol descibed by Das et al. [4]. Briefly the methanolic extract of the beverage was concentrated under vacuum and derivatized using N-methyl-N-(trimethylsilyl) trifluoroacetamide (MSTFA). GC–MS analysis was carried out in a Shimadzu GC 2010 Plus-triple Quadrupole.The peaks were identified using National Institute of Standards and Technology (NIST) library, USA and the column bleeds were removed prior to analyses.

### 1.3 Next Generation Sequencing (NGS) of metagenomic DNA extracted from rice beverage

Metagenomic DNA from rice beverage was extracted following protocol described by Das et. al [12]. The DNA was subjected to NGS with sequencing service provider Macrogen Inc. (Seoul, Republic of Korea). V3-V4 region of bacterial 16S rDNA was amplified using 341F–805R primers. The NGS library was prepared using the Nextera XT library preparation kit following the Illumina MiSeq protocol (Illumina Inc., 2017). Sequencing was carried out in an Illumina MiSeq machine (MiSEq 2500) following 2 × 300 bp paired end chemistry with the multiplexed pooled samples. The trimmed sequences were checked for FASTQ and the high quality regions with an average Phred score higher than 20 considered for further analysis using QIIME (Quantitative Insights Into Microbial Ecology) pipeline. Each sequence was assigned taxonomy at phylum, class, order, family and genus levels with 97% homology.

### 1.4 Animals and experimental design

Five weeks old (36±6 gm) male Swiss albino mice (n=6 per group) were used for the study. Animals were housed in polypropylene cages (Tarsons, India), each cage containing 3 mice. They were fed *ad libitum* standard rodent pellet diet and water. Animals were divided randomly, based on treatments. A 200-230 µl of RB, IF and SF were administered whereas the controls (CT) received sterile water using gavage. The dosage was equivalent to the reported human intake of the beverage based on their body weight to make a consistent gavage volume. All experiments were conducted following the guidelines of the Committee for the Purpose of Control And Supervision of Experiments on Animals (CPCSEA), Ministry of Environment, Forest and Climate Change, Government of India, New Delhi and approved by institutional animal ethical committee (approval no. IASST/IAEC/2016-17/07).

### 1.5 Behavioral analyses

After 30 days of treatment, elevated plus (EP) and Y maze tests were conducted following standard procedures to study anxiety and spatial memory, respectively using methods described earlier [13, 14]. The tests were conducted in an isolated room by placing the mouse in the mazes. An infrared camera (D3D, United Kingdom) was mounted parallel to the mazes to track the animal’s movement.The mazes were made up of acrylic material, matte black in color and cleaned with non-abrasive paper towels using 70% ethanol before and after each test. Animals were acclamatized prior to start of the study.

### 1.6 EP maze test

The EP maze consisted of four arms, 30 cm long and 5 cm wide and was elevated 50 cm above the floor. One pair of opposing arms was exposed with a 3 cm high parameter border along the outer edges while the other pair was enclosed with opaque walls of 12 cm high. The animal was allowed to explore the maze freely for 300 seconds and the process was recorded. Out of 300 seconds, time spent in the open arm was calculated as a measure of anxiolysis [15].

### 1.7 Y maze test

Th Y maze consisted of three arms of equal lengths of 38 cm separated from each other by 120 degrees and designated as A, B and C for scoring. They were habituated to explore the mazes for 10 min having access to two arms. During the test run, a mouse was placed at the centre of the maze and allowed to move freely through the maze during a 5 min session. Entry of hind limb into an arm of the Y maze was considered for the scoring. Spontaneous alternation (SA), defined as the tendency of the animal to explore new arms was calculated as a measure of spatial memory [16].

### 1.8 Statistical analyses

Behavioral parameters on EP and Y maze across the groups were measured and one-way ANOVA was performed using SigmaPlot to determine the differences among the treatment groups. Statistical significance was considered at (*p*<0.05).

## 2 Results and discussion

The biochemical properties of the beverage **(Table 1)** are in line with the previous reports[4]. Out of 13 amino acids detected by GC-MS **(Table 2)**, Methionine, Phenylalanine,Threonine and Valine fell into the category of essential amino acids. This is a good indication in terms of the nutritional content of the beverage. Moreover, the complex, L-Val-L-Leu has been reported to promote healthy gut bacteria [17]. Next, beta Alanine has been effective in reducing anxiety and increasing BDNF expression in rats. Similarly, Phenylalanine and Glutamic acid are also reported to have neuromodulatory effects [18, 19]. Other components, *viz*. Cinnamic acid and Threonic acid have also shown memory enhancing activities [20]. Further, sugars like Allose and Trehalose exert neuroprotective activities [21, 22].

**Table 1.** Biochemical parameters of the rice beverage.

| Parameter | Inferences |
| --- | --- |
| pH | $3.98 \pm 0.005$ |
| Alcohol by volume | $10.55 \pm 0.07$ |
| Reducing sugars | $3.86 \pm 0.33$ |
| Percent inhibition of DPPH free radicals | $29.80 \pm 0.21$ |
| Antioxidant activity (Ascorbic acid microgram/ml) | $1.22 \pm 0.12$ |
| Total phenolic content (Gallic acid equivalent microgram/ml) | $0.392 \pm 0.01$ |

**Table 2.**
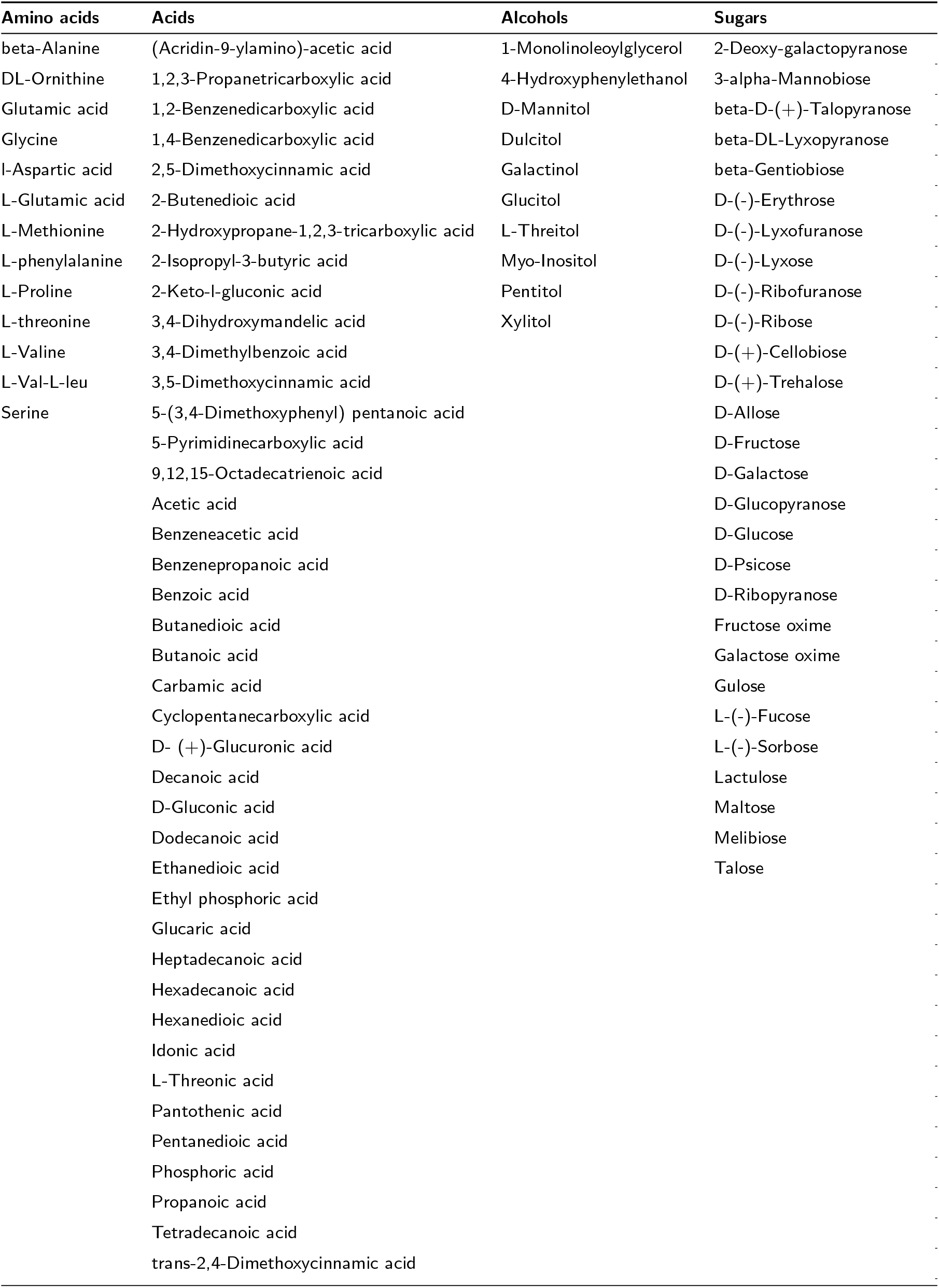
List of metabolites obtained by GC-MS of the rice beverage.

From the NGS data, 574030 quality reads showed a significant homology with predicted rRNA sequences in the Greengene database. A total of 157 OTUs were observed of which *Bacillus* was the most abundant genus (42.5%) **(Figure 3)**. Similar to our findings, *Bacillus* has been found to be the most dominant genus in Chinese rice wine and plays important roles like development of flavor, production of important metabolic products and synthesis of organic acids [23]. Next, an unclassified genus from *Enterobacteriaceae* was dominant, which is also present commonly in wines and other fermented beverages and exert antimicrobial activity by producing bacteriocins [24].Next dominant taxa belonged to *Leuconostocaceae* produces lactic acid from carbohydrate fermentation, and confers probiotic attributes. Moreover, *Lactococcus* and *Lactobacillus* have also been reported to exert such activities.

**Figure 3.**
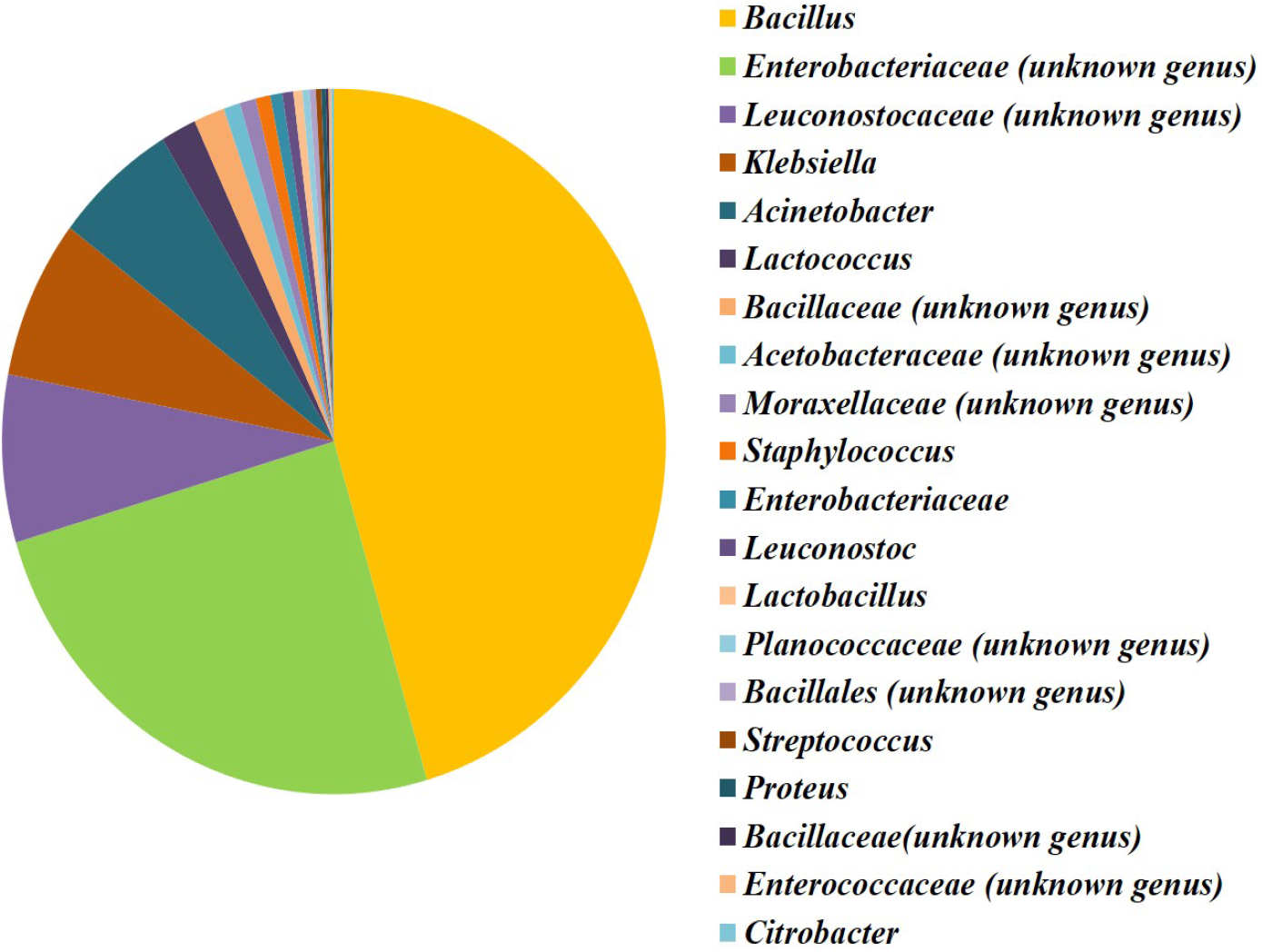
Bacterial diversity of the traditionally fermented rice beverage, based on 16s rDNA sequences. A total of 157 bacterial genera were detected, of which, 20 were present in a relative abundance of at least 0.1%.

In EP maze, more time spent in the open arms reflects less anxiety. IF treatmet spent longest duration (162.55 sec) in the open arm of the EP maze followed by RB (160.3 sec) and SF (87.65 sec) treatments. Significant differences were observed in RB and IF groups compared to the SF group (*p*<0.05)**(Figure 4)**.A higher SA % in Y maze reflects augmented spatial memory. RB showed the highest SA % (74.56 %), followed by IF (73.65 %) and SF (62.75 %). The spatial memory was improved significantly(*p*<0.05) in all treated groups with respect to CT group **(Figure 5)**.

**Figure 4.**
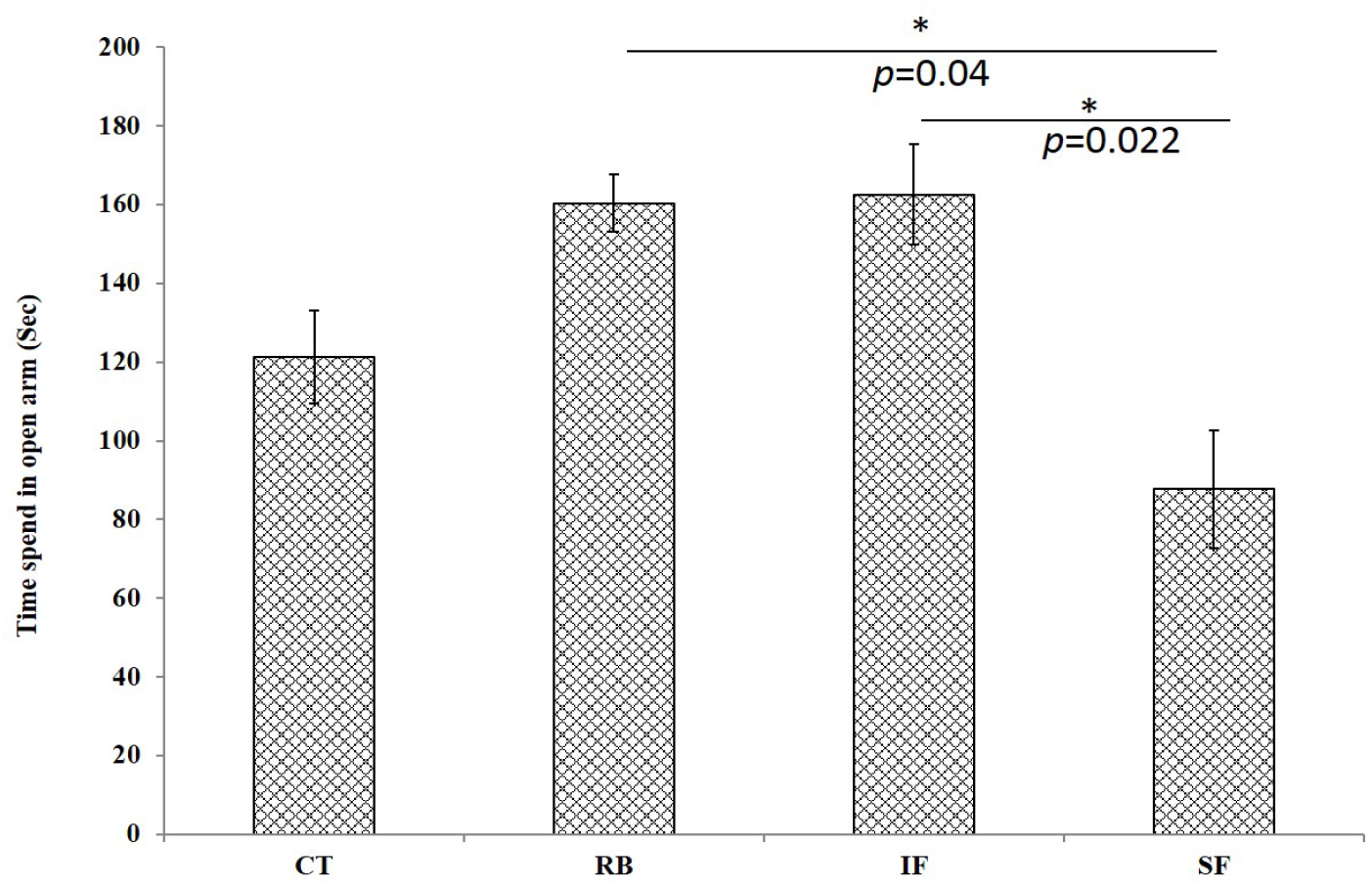
Effects of different rice beverage fractions on behavioral parameters of mice as observed in the EP maze. During a 300 sec test, the time spent on the open arm of the maze (expressed as mean ± SEM) denotes less anxiety like behavior in the animals. *Significant difference (*p*<0.05).

**Figure 5.**
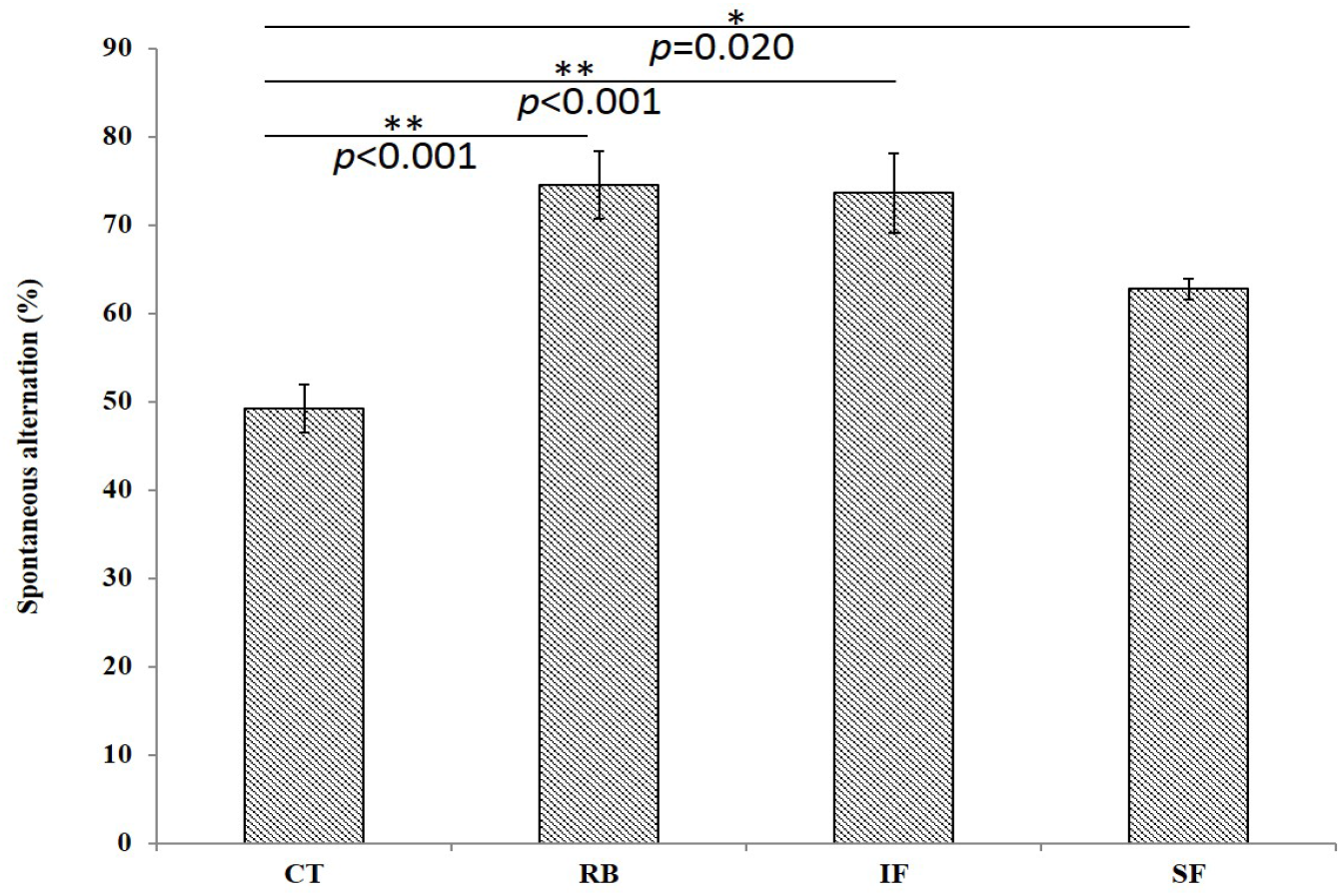
Spontaneous alternation (%) as calculated by scoring the entries into the new arms of the Y maze. The animal’s ability to explore new arms of the maze is related to an improved spatial memory. *Significant difference (*p*<0.05),** (*p*<0.05) *Significant difference (*p*<0.05),** (*p*<0.05)

In the present study, the IF was devoid of alcohol and consisted of microbes and nutrients present in the fermented residues whereas the SF was devoid of microbes but sugars and other dissolved constituents were present. The rice beverage, in contrary was a combination of these two fractions. Presence of prebiotic and prebiotic components in the beverage is an indicative of ”synbiotic” effects, wherein an array of health beneficial properties could be attributed which could include regulation of mental well being. In EP maze the RB and IF fractions showed differences with the SF. This indicates that less anxiety like symptopms resulted due to the effects of bacteria and other nutrients present in both. Moreover, the microbe free and alcohol containing SF treatment showed lowest redcution in anxiety like behavior. Interestingly, a synergistc effect was displayed in Y maze due to the combined effects of different fractions. Though the spatial memory improved in all the treatments compared to CT,such effects were on a lower side in the SF treatment. This could have resulted due to the detrimental effects alcohol, in the soluble fraction which is known to affect the hippocampus adversely. The bidirectional interaction between the central nervous system and the enteric nervous system, known as the gut brain axis (GBA), exhibits the communication between emotional and cognitive centers of the brain with the gastrointestinal tract [25]. Notably, this communication between gut and brain is regulated via the microbes by several pathways including the immune system, the vagus nerve, or by modulation of neuroactive compounds by the microbiota. Therefore, the microbes are considered to be key regulators of this communication, thereby regulating the mental well being by the pshycobiotic activity.With extensive research on GBA, the role of microbiota in regulation of brain, behavior and cognition is becoming clearer[26].Prebiotic components are yet another trait to curb the neurological complications [27].Inclusion of prebiotics and probiotics have been known to palliate psychiatric disorders like panic disorder, generalized anxiety disorder, major depression and impaired brain function due to IBS [28].The dietary components (prebiotics and probiotics) that influence the brain and behavior are defined as psychobiotics which modulate the gut brain axis[29]. As diet remains a major factor to influence the brain and behavior,inclusion of psychobiotic elements can be effective for prevention and treatment of certain neurological diseases [30].

Altogether, it is clearly evident that the components of the beverage affect the behavior and cognition that provides a clue of interference with the gut brain axis. However, further research is required for subsequent validations considering several parameters such as dose and persistence of such effects. These studies would provide novel insights for the therapeutic potential of rice beverage for neurological complications.

## 3 Conclusion

Administration of rice beverage fractions showed improved brain functions in mice. This suggests that the prebiotic and probiotic components of rice beverage may be responsible for such behavior. Whether these results could be considered for the neuromodulatory effects of the beverage, further studies could target on its utilization as a functional food based therapeutic interventions.

## Acknowledgements

This study was supported by the core fund of IASST and Department of Biotechnology (DBT, Govt. of India) funded unit of excellence project (BT/550/NE/U-EXCEL/2014). The research was carried out in the Institutional Level Biotech Hub of IASST. Authors are thankful to Mrs. Anima Baishya for sharing traditional knowledge on rice beverage preparation. Mr. Milan Jyoti Das, Mr. Gwhwm Basumatary and Mr. Abinash Nath for assistance in the animal rearing and handling.

## Funding

This study was supported by the core fund of IASST and Department of Biotechnology (DBT, Govt. of India) funded unit of excellence project (BT/550/NE/U-EXCEL/2014).

### Abbreviations

ANOVA: Analysis of Variance
DPPH: 2,2-diphenyl-1-picrylhydrazyl
EP: Elevated Plus
GBA: Gut Brain Axis
GC-MS: Gas chromatography–mass spectrometry
MSTFA: N-methyl-N-(trimethylsilyl) trifluoroacetamide
NIST: National Institute of Standards and Technology
NGS: Next Generation Sequencing
OTU: Operational Taxonomic Unit
QIIME: Quantitative Insights Into Microbial Ecology
SA: Spontaneous Alternation
SCFA: Short Chain Fatty Acids

## Ethics approval and consent to participate

All experiments were conducted following the guidelines of the Committee for the Purpose of Control And Supervision of Experiments on Animals (CPCSEA), Ministry of Environment, Forest and Climate Change, Government of India, New Delhi and approved by institutional animal ethical committee (approval no. IASST/IAEC/2016-17/07).

## Authors’ contributions

BB, AA and MRK designed the experiments, BB and AA performed the experiments. Manuscript was drafted by BB and revised by MRK.

## Author details

Molecular Biology and Microbial Biotechnology Laboratory, Institute of Advanced Study in Science and Technolgy, Vigyan Path, Paschim Boragaon, Guwahati, India.

## References

1. Das, A., Deka, S.: Fermented foods and beverages of the north-east india. International Food Research Journal 19(2), 377 (2012)

2. Bora, S.S., Keot, J., Das, S., Sarma, K., Barooah, M.: Metagenomics analysis of microbial communities associated with a traditional rice wine starter culture (xaj-pitha) of assam, india. 3 Biotech 6(2), 153 (2016)

3. Das, A., Deka, S., Miyaji, T.: Methodology of rice beer preparation and various plant materials used in starter culture preparation by some tribal communities of north-east india: A survey. International Food Research Journal 19(1), 101 (2012)

4. Das, A.J., Khawas, P., Miyaji, T., Deka, S.C.: Hplc and gc-ms analyses of organic acids, carbohydrates, amino acids and volatile aromatic compounds in some varieties of rice beer from northeast india. Journal of the Institute of Brewing 120(3), 244–252 (2014)

5. Korneeva, O., Cheremushkina, I., Glushchenko, A., Mikhaĭlova, N., Baturo, A., Romanenko, É., Zlygostev, S.: Prebiotic properties of mannose and its effect on specific resistance. Zhurnal mikrobiologii, epidemiologii, i immunobiologii (5), 67–70 (2012)

6. Wongputtisin, P., Ramaraj, R., Unpaprom, Y., Kawaree, R., Pongtrakul, N.: Raffinose family oligosaccharides in seed of glycine max cv. chiang mai60 and potential source of prebiotic substances. International Journal of Food Science & Technology 50(8), 1750–1756 (2015)

7. Khan, M.R., Bhaskar, B., Adak, A., Talukdar, N.C.: Rice Based Beverage with High Alcohol Content and Method Therefore. 201731006470, (2017)

8. Caputi, A., Ueda, M., Brown, T.: Spectrophotometric determination of ethanol in wine. American Journal of Enology and Viticulture 19(3), 160–165 (1968)

9. Miller, G.L.: Use of dinitrosalicylic acid reagent for determination of reducing sugar. Analytical chemistry 31(3), 426–428 (1959)

10. Leong, L., Shui, G.: An investigation of antioxidant capacity of fruits in singapore markets. Food chemistry 76(1), 69–75 (2002)

11. Singleton, V.L., Orthofer, R., Lamuela-Raventús, R.M.: [14] analysis of total phenols and other oxidation substrates and antioxidants by means of folin-ciocalteu reagent. Methods in enzymology 299, 152–178 (1999)

12. Das, S., Deb, D., Adak, A., Khan, M.R.: Exploring the microbiota and metabolites of traditional rice beer varieties of assam and their functionalities. 3 Biotech 9(5), 174 (2019)

13. Komada, M., Takao, K., Miyakawa, T.: Elevated plus maze for mice. JoVE (Journal of Visualized Experiments) (22), 1088 (2008)

14. Miedel, C.J., Patton, J.M., Miedel, A.N., Miedel, E.S., Levenson, J.M.: Assessment of spontaneous alternation, novel object recognition and limb clasping in transgenic mouse models of amyloid-β and tau neuropathology. JoVE (Journal of Visualized Experiments) (123), 55523 (2017)

15. Pellow, S., Chopin, P., File, S.E., Briley, M.: Validation of open: closed arm entries in an elevated plus-maze as a measure of anxiety in the rat. Journal of neuroscience methods 14(3), 149–167 (1985)

16. Nakagawasai, O., Onogi, H., Mitazaki, S., Sato, A., Watanabe, K., Saito, H., Murai, S., Nakaya, K., Murakami, M., Takahashi, E., et al.: Behavioral and neurochemical characterization of mice deficient in the n-type ca2+ channel α1b subunit. Behavioural brain research 208(1), 224–230 (2010)

17. Hatanaka, M., Morita, H., Aoyagi, Y., Sasaki, K., Sasaki, D., Kondo, A., Nakamura, T.: Effective bifidogenic growth factors cyclo-val-leu and cyclo-val-ile produced by bacillus subtilis c-3102 in the human colonic microbiota model. Scientific reports 10(1), 1–9 (2020)

18. Lou, H.: Dopamine precursors and brain function in phenylalanine hydroxylase deficiency. Acta Paediatrica 83, 86–88 (1994)

19. Zhou, Y., Danbolt, N.C.: Glutamate as a neurotransmitter in the healthy brain. Journal of neural transmission 121(8), 799–817 (2014)

20. Surman, C., Vaudreuil, C., Boland, H., Rhodewalt, L., DiSalvo, M., Biederman, J.: L-threonic acid magnesium salt supplementation in adhd: an open-label pilot study. Journal of dietary supplements, 1–13 (2020)

21. Huang, T., Gao, D., Hei, Y., Zhang, X., Chen, X., Fei, Z.: D-allose protects the blood brain barrier through pparγ-mediated anti-inflammatory pathway in the mice model of ischemia reperfusion injury. Brain research 1642, 478–486 (2016)

22. Khalifeh, M., Read, M.I., Barreto, G.E., Sahebkar, A.: Trehalose against alzheimer’s disease: Insights into a potential therapy. BioEssays 42(8), 1900195 (2020)

23. Xu, J., Wu, H., Wang, Z., Zheng, F., Lu, X., Li, Z., Ren, Q.: Microbial dynamics and metabolite changes in chinese rice wine fermentation from sorghum with different tannin content. Scientific reports 8(1), 1–11 (2018)

24. Zhang, J., Zhu, X., Xu, R., Gao, Q., Wang, D., Zhang, Y.: Isolation and identification of histamine-producing enterobacteriaceae from qu fermentation starter for chinese rice wine brewing. International journal of food microbiology 281, 1–9 (2018)

25. Martin, C.R., Mayer, E.A.: Gut-brain axis and behavior. In: Intestinal Microbiome: Functional Aspects in Health and Disease vol. 88, pp. 45–54. Karger Publishers, ??? (2017)

26. Cryan, J.F., Dinan, T.G.: Mind-altering microorganisms: the impact of the gut microbiota on brain and behaviour. Nature reviews neuroscience 13(10), 701–712 (2012)

27. Shah, N.P.: Functional cultures and health benefits. International dairy journal 17(11), 1262–1277 (2007)

28. Pusceddu, M.M., Murray, K., Gareau, M.G.: Targeting the microbiota, from irritable bowel syndrome to mood disorders: focus on probiotics and prebiotics. Current pathobiology reports 6(1), 1–13 (2018)

29. Sarkar, A., Lehto, S.M., Harty, S., Dinan, T.G., Cryan, J.F., Burnet, P.W.: Psychobiotics and the manipulation of bacteria–gut–brain signals. Trends in neurosciences 39(11), 763–781 (2016)

30. Luna, R.A., Foster, J.A.: Gut brain axis: diet microbiota interactions and implications for modulation of anxiety and depression. Current opinion in biotechnology 32, 35–41 (2015)

